# Is adaptive foraging adaptive? A resource-consumer eco-evolutionary model

**DOI:** 10.1101/2023.03.22.533765

**Authors:** Leo Ledru, Jimmy Garnier, Océane Guillot, Erwan Faou, Camille Noûs, Sébastien Ibanez

**Affiliations:** Univ. Savoie Mont Blanc, Univ. Grenoble Alpes, CNRS, UMR 5553 LECA, France; CNRS, Univ. Savoie Mont Blanc, UMR 8050 LAMA, France; INRIA, Universite Rennes 1, UMR CNRS 6625 IRMAR, France; Laboratory Cogitamus

**Keywords:** phenotypic plasticity, adaptive foraging, eco-evolutionnary dynamics, community stability

## Abstract

Phenotypic plasticity has important ecological and evolutionary consequences. In particular, behavioural phenotypic plasticity such as adaptive foraging (AF) by consumers, may enhance community stability. Yet little is known about the ecological conditions that favor the evolution of AF, and how the evolutionary dynamics of AF may modulate its effects on community stability. In order to address these questions, we constructed an eco-evolutionary model in which resource and consumer niche traits underwent evolutionary diversification. Consumers could either forage randomly, only as a function of resources abundance, or adaptatively, as a function of resource abundance, suitability and consumption by competitors. AF evolved when the niche breadth of consumers with respect to resource use was large enough and when the ecological conditions allowed substantial functional diversification. In turn, AF promoted further diversification of the niche traits in both guilds. This suggests that phenotypic plasticity can influence the evolutionary dynamics at the community-level. Faced with a sudden environmental change, AF promoted community stability directly and also indirectly through its effects on functional diversity. However, other disturbances such as persistent environmental change and increases in mortality, caused the evolutionary regression of the AF behaviour, due to its costs. The causal relationships between AF, community stability and diversity are therefore intricate, and their outcome depends on the nature of the environmental disturbance, in contrast to simpler models claiming a direct positive relationship between AF and stability.

## 1 Introduction

Phenotypic plasticity has become central to evolutionary theory (West-Eberhard, 2003; Pfennig, 2021) as it may mitigate environmental changes (Chevin et al., 2013; Vedder et al., 2013; Charmantier et al., 2008). Phenotypic plasticity commonly occurs when a variety of resources are available to consumers investing more or less time on each resource according to its suitability. The resulting *relative foraging efforts* (sensu Abrams, 2010) depend on the (mis)match between the defensive and counter-defensive traits (e.g. Clissold et al., 2009), and the nutritional quality of the resources and the requirements of the consumers (e.g. Behmer and Joern, 2008). Relative foraging efforts sometimes correspond to the best compromise between suitability and abundance, an outcome called *optimal foraging* (MacArthur and Pianka, 1966; Loeuille, 2010). However optimal foraging might be difficult to achieve when the identity and abundance of resources vary over time and space, because foraging optimization is not instantaneous (Abrams, 1992, 2010). Under such circumstances, consumers may nevertheless redirect their relative foraging efforts towards more profitable resources in order to increase their energy intake. The ability to adjust relative foraging efforts is a type of behavioural plasticity called *adaptive foraging* (AF, Valdovinos et al., 2013).

Phenotypic plasticity often results from evolution by natural selection (Nussey et al., 2005; Peluc et al., 2008; Van Kleunen and Fischer, 2001), but not always, especially in the context of environmental changes (Merilä and Hendry, 2014). The extent to which phenotypic plasticity is adaptive has been underexplored in the context of AF because previous theoretical works ignored the evolutionary dynamics of AF, focusing instead on food-web stability (Kondoh, 2003; Uchida and Drossel, 2007; Heckmann et al., 2012) or food web structure (Beckerman et al., 2006). Abrams (2003) modelled the evolution of the general foraging effort, corresponding to the overall amount of time and energy invested in foraging (e.g. Dill, 1983), in function of the trade-off with predation risk. *General* foraging effort differs from AF, that in contrast focuses on the adjustment of *relative* foraging efforts, i.e. how the general foraging effort is distributed across the different resources. Although the AF strategy tends to increase fitness, in some situations AF may reduce it by increasing predation risk (Abrams, 2003; Pangle et al., 2012; Wang et al., 2013; McArthur et al., 2014; Costa et al., 2019), preventing efficient thermoregulation (du Plessis et al., 2012; Van de Ven et al., 2019) or increasing searching time for resources (Randolph and Cameron, 2001; Bergman et al., 2001; Fortin et al., 2004). Since AF faces several trade-offs with life-history components, its evolution should depend on ecological parameters such as mortality rate, resource searching time or consumer niche width.

The first aim of the present study is therefore to understand, using a theoretical model, under which ecological conditions the ability of consumers to forage adaptatively is subject to evolution by natural selection. In other words: is adaptive foraging itself adaptive? We define AF as a change in relative foraging efforts that directly increases *energy intake*, but not necessarily *fitness*, in contrast with Loeuille (2010) who defined AF as “changes in resource or patch exploitation by consumers that give the consumer a higher fitness compared with conspecifics that exhibit alternative strategies”. Our restricted definition is justified by the need to explore how the trade-off between energy intake and other life-history components modulates the evolution of AF. Moreover, consumers are affected by environmental changes, either directly (Bale et al., 2002; Staley and Johnson, 2008; Scherber et al., 2013) or indirectly through changes affecting their resources. For instance, environmental changes may induce a shift in resource phenology (Altermatt, 2010; Kerby et al., 2012; Portalier et al.) or alter resource chemistry (Bidart-Bouzat and Imeh-Nathaniel, 2008; Rasmann and Pellissier, 2015). As a result, the diet preferences of consumers may be altered (Rasmann et al., 2014; Rosenblatt and Schmitz, 2016; Boersma et al., 2016), suggesting that environmental disturbances should lead to the evolution of AF. However as disturbances may also reduce the functional diversity of available resources (Thuiller et al., 2006; Buisson et al., 2013), the evolutionary response of the AF strategy to environmental changes is unclear.

Although phenotypic plasticity generally results from evolution by natural selection, as outlined above, it also generates evolutionary changes (Simpson, 1953; Baldwin, 1896; Laland et al., 2014), with genes acting as followers (West-Eberhard, 2003). In the context of AF, the consumption of novel or unusual resources through behavioral plasticity might trigger subsequent adaptations that favour the use of these resources. This would increase the diversity of the traits involved in resource use, such as counter-defences and nutritional requirements. The second motivation is therefore to investigate how AF can alter the evolution of these consumer traits, as well as those of their resources (defenses, nutritional quality). In particular, we expect AF to affect the functional diversity of consumers and resources, through its effects on diet breadth.

The evolutionary dynamics of phenotypic plasticity has important ecological consequences (Miner et al., 2005; Turcotte and Levine, 2016), which in turn can feed back into the evolutionary dynamics. In the case of AF, behavioural plasticity in diet choice can favour the persistence of consumers in unusual environments and rescue them in the face of environmental changes (e.g. Varner and Dearing, 2014; Kowalczyk et al., 2019). Previous theoretical studies have indeed shown that AF promotes community stability (Křivan and Schmitz, 2003; Abrams and Matsuda, 2004; Kondoh, 2003; Uchida and Drossel, 2007). The third motivation is to test if this positive relationship holds when both AF and the functional traits of consumers and resources are subject to evolutionary dynamics. In this eco-evolutionary context, it is uncertain whether the evolution of AF stabilises communities directly or indirectly, through its effects on functional diversity.

The main questions outlined earlier are sketched in Figure 1:

**Figure 1:**
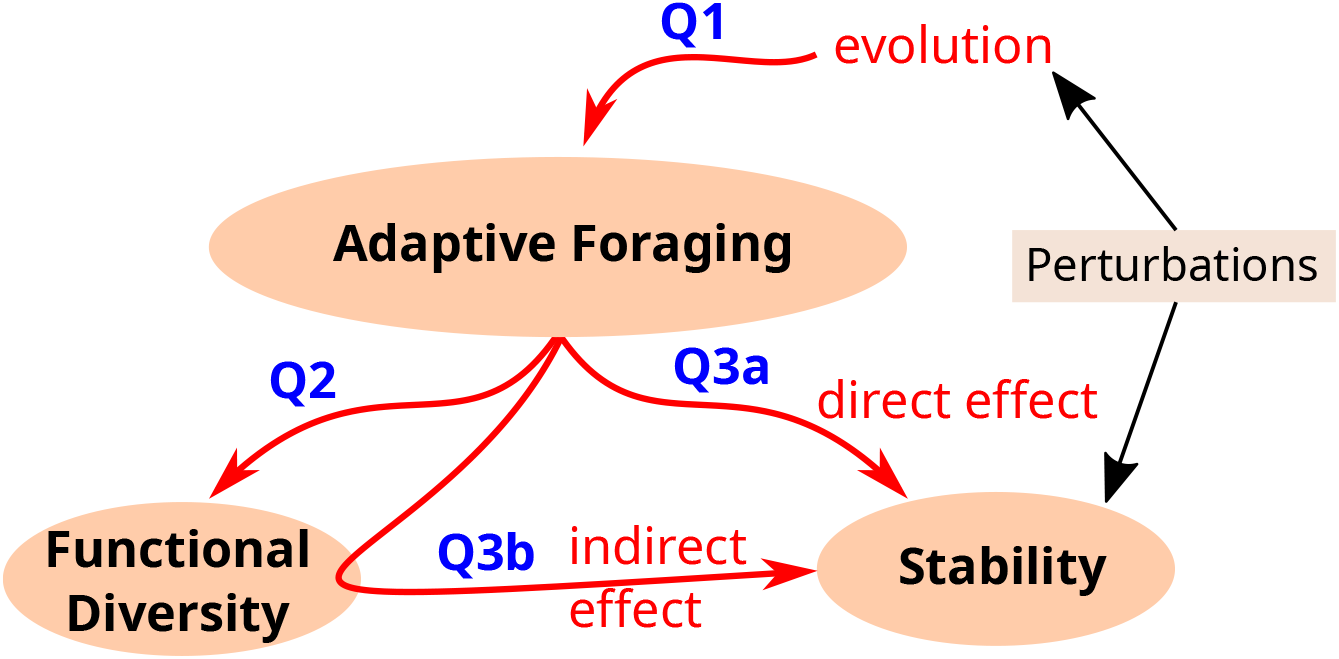
Overview of the main questions: (Q1) Under which ecological conditions does AF evolve? (Q2) Does the evolution of AF increases the diversity of traits involved in the resource-consumer interaction? (Q3) Does the evolution of AF enhances the stability of the resource-consumer system, either directly (Q3a) or through its effects on functional diversity (Q3b)?

- Question 1. Under which ecological conditions is AF evolutionary adaptive? How do environmental disturbances alter the evolution of AF?
- Question 2. When AF evolves, what are its effects on the diversity of the traits involved in the resource-consumer interaction?
- Question 3. What is the effect of the evolution of AF on the stability of the resource-consumer system? Are these effects direct (Q3a) or indirect, mediated by the influence of AF on functional diversity (Q3b)?

To address these issues, we build an eco-evolutionary model in which a consumer species feeds on a resource species. Both species are characterized by an ecological trait; the resource is the most suitable for the consumer when both traits match. In addition, the consumers carry a foraging trait measuring the extent to which they select the resources allowing the largest intake, or instead forage randomly and consume the resources as a function of their abundance. Ecological and foraging traits are subject to evolution; starting from monomorphic initial conditions, they rapidly diversify and reach a stationary regime characterized by a stable diversity of ecological and foraging traits. The stationary regime is then subjected to various environmental disturbances, to test how the evolution of AF responds to environmental changes, and how this cascades down on the ecological properties of the resource-consumer system.

### 2 Model and methods

#### 2.1 Aresource-consumer niche model

An eco-evolutionary model is developed to describe the dynamics of a consumer population feeding, with various individual foraging strategies, on a resource population. Consumers compete for resources both directly and indirectly. Individuals are characterized by quantitative traits: the niche traits *x* and *y* of consumers and resources, respectively, and the adaptive foraging trait *z* of consumers. The niche traits affect competition between individuals as well as interactions between consumer and resource individuals. The foraging trait *z* affects the foraging strategy of the consumers through their foraging efforts *φ*. The model describes the time dynamics of the trait densities of resources *r*(*t, y*) and consumers *c*(*t, x, z*); the components of the model are detailed in the following sections.

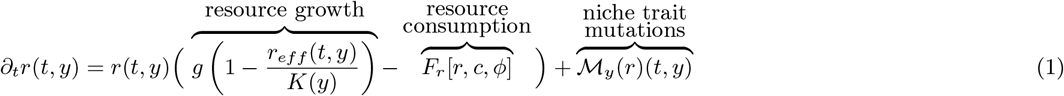

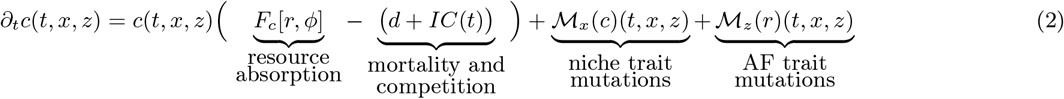

#### Resource-consumer interactions and niche traits

In the absence of consumers, resources grow logistically with an intrinsic rate *g*, independent from the niche trait *y*. Competition between resources depends on the niche trait *y* through the carrying capacity *K*(*y*) of individuals with trait *y* and *r*_*eff*_ (*t, y*), the effective population density perceived by an individual with trait *y* at time *t*. The effective density depends on the phenotype distribution of the population and the competition strength *K*_*eff*_ (*y −y*^*t*^) exerted by an individual with trait *y*^*t*^ on an individual with trait *y*:

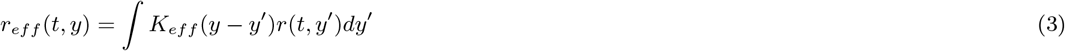

The functions *K* and *K*_*eff*_ are normally distributed around *y* = 0 with variances *σ*_*K*_ and *σ*_*C*_ respectively (Table A1 and Fig. A1). In the presence of consumers, resources are exploited at rate *F*_*r*_ [*r, c, φ*], whereas the consumer density increases through resource absorption at a rate *F*_*c*_[*c, φ*]. Although these rates vary with the consumers foraging efforts *φ*, they crucially depend on the effective interaction strength ∆(*x− y*) between consumer and resource individuals. The function ∆ is normally distributed around 0 with a variance *σ*, which measures the extend to which consumers can deal with a variety of resource types (Table A1). The variance parameter *σ* is chosen similarly to previous models (see e.g. Dieckmann and Doebeli, 1999; Egas et al., 2005), but it is not subject to evolution as in Egas et al. (2005). The interactions are described by a Holling type II functional response, which provides the following consumption and absorption rates:

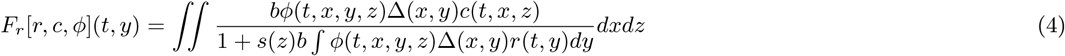

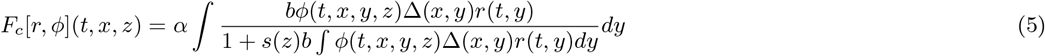

with *α* the conversion coefficient, *b* the extraction coefficient and *s*(*z*) the searching time, which depends on the foraging trait *z* as explained below. Moreover, consumer density is affected by mortality at a constant rate *d* and by direct competition at a rate *I C* where *C*(*t*) = *c*(*t, x, z*)*dxdz* is the total biomass of consumer and *I* the intraspecific competition between consumers for other limiting factors than resources.

#### Foraging strategies and adaptive foraging trait

Consumers can use two different foraging strategies during their foraging time: Random Foraging (RF) or Adaptive Foraging (AF). When using RF, the consumer randomly forages its environment without selecting resources. The resulting efforts *φ*_*RF*_ is proportional to the density of the resources:

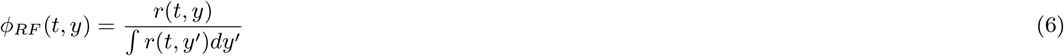

Conversely, when using AF, consumers actively search for resources, that maximize their energy intake, that depends on the resource availability and suitability (e.g. Sundell et al., 2003). The resulting relative foraging efforts *φ*_*AF*_ may change over time as follows:

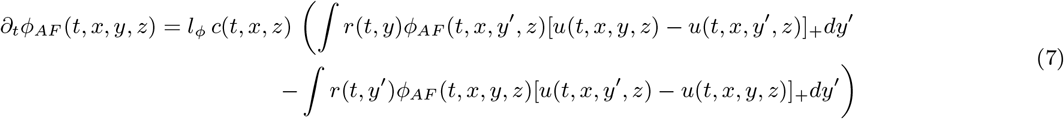

The quantity *φ*_*AF*_ is analogous to the behavioral trait *z* in Abrams and Matsuda (2004). The potential gain *u*(*t, x, y, z*) of a consumer with traits (*x, z*) on a resource with trait *y* depends on its foraging efforts as well as the resource suitability and availability:

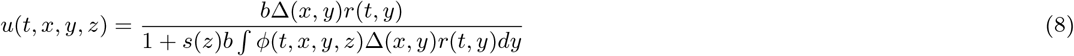

The AF dynamics allow consumers to compare the benefits received from different resources. As a result, consumers increase their efforts on the most beneficial resources and reduce them on sub-optimal resources. The comparison of resources is assumed time consuming, the efforts are therefore not adjusted instantaneously but exponentially fast at a rate *l*_*φ*_. When the adjustment rate *l*_*φ*_ becomes large, the adaptive foraging strategy becomes closer to the optimal foraging strategy maximizing the potential gain *u* (MacArthur and Pianka, 1966; Loeuille, 2010). Moreover, the searching time *s*(*z*) also increases with the foraging trait: *s*(*z*) = *s*_*min*_ + *z*(*s*_*max*_ *−s*_*min*_) (Figure A1d). This relationship introduces a trade-off between the AF strategy and the searching time.

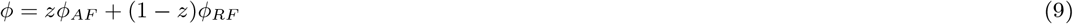

Finally, the effective consumer foraging strategy depends on its foraging trait *z ∈* [0, 1], which corresponds to the proportion of its general foraging effort spent using the AF strategy. The effective consumer efforts is thus:

#### Mutation of traits and diffusion approximation

Due to mutations, the niche traits and the foraging trait can evolve independently. Foraging behaviour can indeed be heritable in nature (Wallin, 1988; Lemon, 1993). Since ecological and evolutionary dynamics occur on the same time scale, mutants are constantly introduced through the diffusion of traits:

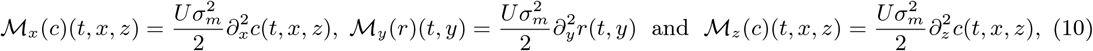

where *U* is the mutation frequency and 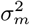 is the variance of the mutational effects. This approach contrasts with the adaptive dynamic framework, in which a mutant phenotype is introduced sequentially and persists only if its invasive fitness is positive (Geritz et al., 1998).

### 2.2 Analysis of the model

#### Sensitivity analysis on the mean foraging trait

The model is investigated numerically using MATLAB (code available on GitHub https://github.com/leoledru/Adaptive-Foraging). The niche traits are discretized into 31 equally distanced values (11 values for the foraging trait). In the simulations, when the density of a resource or a consumer phenotype drops below the critical threshold *ε* = 10^*−*4^, the density is set to 0 to save computational time. The simulations start with monomorphic populations at the niche center (*y* = *x* = 0) and consumers have a purely random foraging strategy (*z* = 0). To investigate the ecological conditions leading to the evolution of AF, a global sensitivity analysis is performed using Partial Rank Correlations Coefficients (PRCC, Saltelli et al., 2004), on the mean foraging trait value of the consumer population 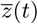 and the tested parameter defined by:

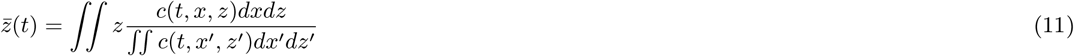

The analysis focuses on the parameters *σ, σ*_*K*_, *s*_*max*_, *d, I, g* (Table 1) with 5000 parameter sets sampled in their ranges.

**Table 1:** Parameters of the model with their reference values used for the analysis of the response to disturbances, and the range used for the 6 parameters tested by the sensitivity analysis. The last column corresponds to the PRCC values, that is the correlation between the mean foraging trait 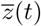 and the tested parameter

| Parameters |  | Values for the<br>response to disturbances | Ranges for the<br>sensitivity analysis | PRCC<br>values |
| --- | --- | --- | --- | --- |
| $\sigma$ | Consumers niche width | 0.9 | [0; 1] | 0.28 |
| $\sigma_K$ | Resources niche width | 2.5 | [1; 4] | 0.38 |
| $s_{max}$ | Cost of AF : maximal increase of<br>searching time due to AF | 0.55 | [0.1; 2] | - 0.64 |
| $d$ | Consumers mortality | 0.1 | [0.1; 0.6] | 0.13 |
| $I$ | Competition between consumers<br>(other than for resources) | 0.01 | [0.01; 0.1] | 0.13 |
| $g$ | Rate of resource growth | 0.8 | [0.2; 1.6] | 0.11 |
| $K_0$ | Maximal carrying capacity | 50 | Fixed | |
| $\sigma_C$ | Width of the competition kernel | $\sigma_K - 1$ | Fixed | |
| $\alpha$ | Biomass conversion coefficient<br>from resources to consumers | 0.3 | Fixed | |
| $b$ | Biomass extraction coefficient | 0.5 | Fixed | |
| $l_\phi$ | Rate of change in foraging efforts | 0.5 | Fixed | |
| $s_{min}$ | Cost of AF : minimal increase of<br>searching time due to AF | 0.1 | Fixed | |
| $U$ | Mutation frequency | 0.1 | Fixed | |
| $\sigma_m^2$ | Mean effect of mutation | 0.02 | Fixed | |
| $\varepsilon$ | Extinction threshold | $10^{-4}$ | Fixed | |
| $T$ | Simulation time | 1000 | Fixed | |

#### Effect of AF evolution on biomass, functional diversity, productivity and niche overlap

To assess the effect of AF on the resource-consumer system, several characteristics were measured: the biomass of resources and consumers, their functional dispersion *FDis* (Laliberté and Legendre, 2010), the productivity *Prod*, the niche overlap between consumers *ρ* (Chesson and Kuang, 2008) and the functional match between consumers and their resources. The biomass of resources *R* and consumers *C* are given respectively by *R*(*t*) = ∫ *r*(*t, y*)*dy* ∫ ∫*c*(*t, x, z*)*dxdz*. The functional dispersion *FDis* represents for each population the average absolute deviation from the mean niche trait:

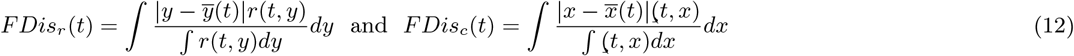

where 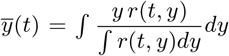 and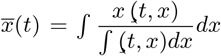 are the mean traits of the resource and consumer and (*t, x*) = ∫*c*(*t, x, z*)*dz* is the biomass of individuals carrying the trait *x* in the consumers population. Productivity corresponds to the net production of biomass by consumers following resource absorption, measured once the system has reached a stationary regime (e.g. Loreau and Hector, 2001; Poisot et al., 2013):

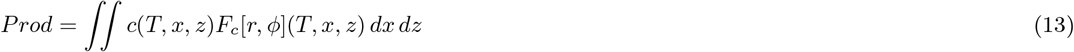

*T* is the time to reach the stationary regime, *T* = 1000 in the simulations below. The niche overlap between two consumers with niche traits *x*_*i*_ and *x*_*j*_ and foraging traits *z*_*i*_ and *z*_*j*_ is defined by the correlation coefficient *ρ*_*ij*_ of their resource absorption:

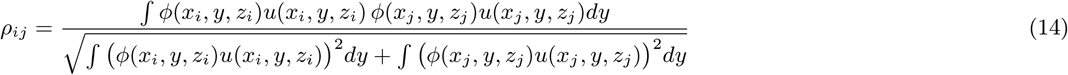

The overall niche overlap between consumers *ρ* is the average of this correlation coefficient of all consumers. The functional match *FM* corresponds to the mean difference between the niche trait of the consumer and the mean niche trait of its diet, that is the resources absorbed by the consumer:

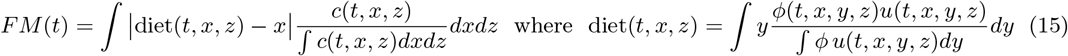

To assess the effects of the evolution of AF on the system, we compare the total biomass *C* of consumers in two situations: a freely evolving AF trait *z* and a fixed RF strategy (*z* = 0). In both cases, the ecological niche traits *x* and *y* are subject to evolution. The communities evolved during 1000 time steps, which is enough time for the system to reach a stationary regime with stable community-level characteristics (A.2). The same comparison was done for all the other system-level characteristics.

#### Effects of environmental disturbance

To understand whether the evolution of AF can rescue consumers from environmental changes, three specific disturbances are considered: (a) an ecosystem disturbance where consumer mortality *d* increases gradually by ∆*d*, (b) a constantly changing environment, where the niche center is shifted at constant speed *c* and (c) an sudden environmental change where the center of the resource niche is instantaneously shifted at a distance ∆*y* from the initial niche center (e.g. Domínguez-García et al., 2019). The mutation process driving the diversification of resources and consumers in the system should help to recover trait diversity after a disturbance. To assess the effects of those disturbances on the resource-consumer system, the proportion of consumer biomass lost after the disturbance is calculated once a new equilibrium is reached. The difference in the mean foraging trait before and after each disturbance is also measured.

The resource-consumer system is initialized with consumers carrying a high mean AF trait 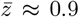with parameter values set as in Table 1). For each disturbance strength and type, the stability metrics of the system with AF evolution is compared to those of the system with RF only, in which the foraging trait of consumers was monomorphic (*z* = 0) and fixed (*M*_*z*_ (*c*) = 0). For all disturbance types, the disturbance strength is increased until the consumer population goes to extinction, in order to compute the maximal disturbance level that the system can tolerate.

## 3 Results

### 3.1 The evolution of adaptive foraging

A typical outcome of the model was the diversification of the resources and consumers along the ecological gradient (Figure 2a). Although the distribution of the consumer foraging trait reached a unimodal distribution (Figure 2a), the consumers positioned at the niche center foraged randomly, while those at the niche edges foraged adaptatively (Figure 2b). In addition, the distributions of the niche traits reached a stationary regime that varied over time due to the AF strategy (Appendix A.2). However, the macroscopic characteristics (functional dispersion, total biomass, productivity, niche overlap and average foraging behavior) stabilized around a steady state; these characteristics will therefore be used to assess the effect of AF evolution on the resource-consumer system.

**Figure 2:**
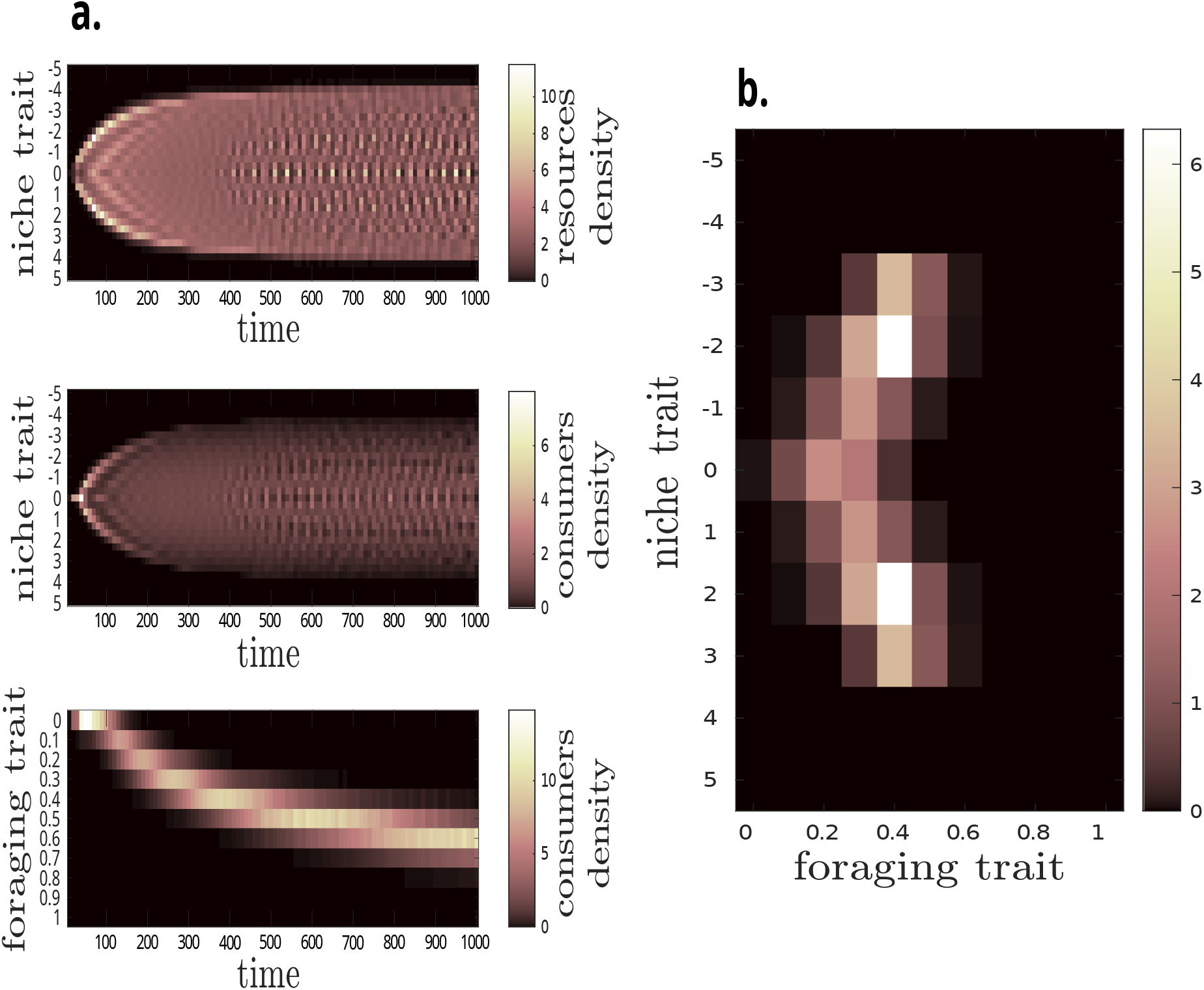
a) Diversification of niche and foraging traits starting from a single resource and consumer at the niche centre, and a RF consumer strategy. Top panel: resource densities *r*(*t, y*). Middle panel: consumer densities ∫*c*(*t, x, z*)*dz*. Bottom panel: foraging trait ∫*c*(*t, x, z*)*dx*. b) The trait distribution of consumers at steady state (1000 time steps).

The PRCC analysis revealed that the six tested parameters played a significant role in the evolution of AF (Table 1 last column). The conditions favouring the evolution of AF (measured by 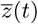 were essentially the following: a consumers ability to exploit a wide range of resources (large *σ*, correlation coefficient 0.28), a wide niche for resources (large *σ*_*K*_, correlation coefficient 0.38), a weak trade-off between AF and searching time (small *s*_*max*_, correlation coefficient *−*0.64), a high consumer mortality *d* (correlation coefficient 0.13), a strong competition between consumers (large *I*, correlation coefficient 0.13) and a high resource growth (large *g*, correlation coefficient 0.11).

### The effects of AF on functional diversity and other macroscopic characteristics

When the evolution of AF produces consumer populations with a high mean foraging trait 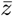, the resource biomass is reduced (e.g. -50% when 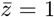) while the consumer biomass increases by 25% on average (Figure 3a). However, the variabililty of the consumer biomass among simulations also increases with 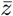. This pattern has also been observed when the foraging trait *z* of a monomorphic population without AF evolution is increased (Figure A3a). Turning to diversity, the evolution of AF increases functional dispersion of both resources and consumers (Figure 3b). Again, when the average foraging trait value was large the consequences on diversity indices become heterogeneous, but this time the effect of AF was almost always positive. The relationship with productivity (i.e the flow of biomass from resources to consumers) was non-linear (Figure 3c). When the system with AF evolution had a rather low mean foraging trait 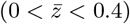productivity increased in comparison to the system without AF. However, when *z* was above 0.4, the productivity gain became smaller and even vanished when 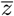 equalled 1. Strong AF also increased the variability of productivity; among the systems with strong AF some had large gains of productivity and others large deficits. Finally, the evolution of AF also decreased the niche overlap between consumers by about 40% when the mean foraging trait was close to 1 (Figure 3d), and increased the functional match between the niche trait of consumers and the mean niche trait of their resources (Figure A4).

**Figure 3:**
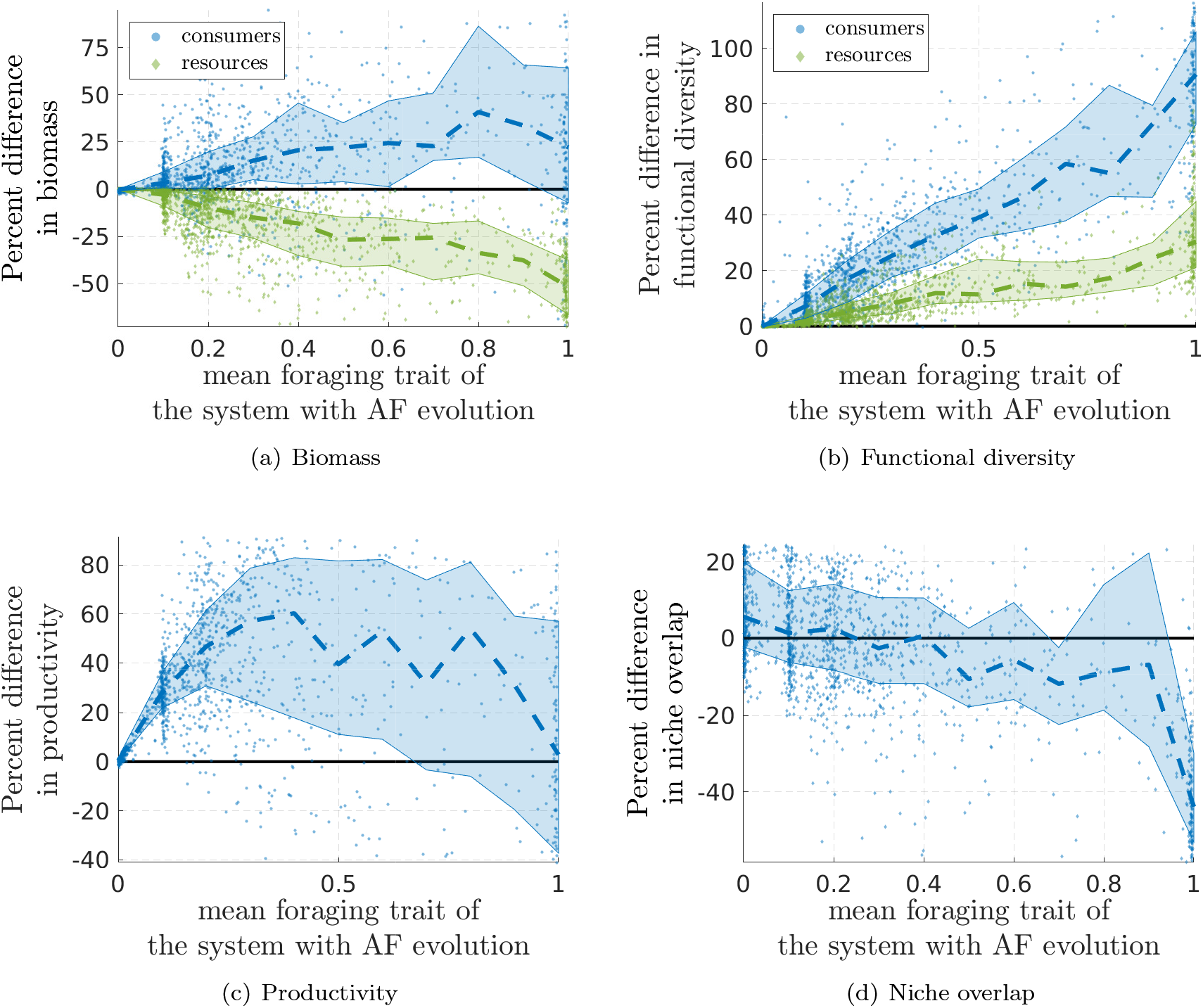
Difference (in %) between systems with AF evolution and fixed RF, for (a) biomass, (b) functional dispersion, (c) productivity, and (d) niche overlap. For each panel, 1500 simulations of 1000 time steps with AF evolution were compared to simulations with fixed RF, the parameters being randomly sampled in the ranges specified in Table 1. Dashed lines: median; areas: 75% confidence intervals.

### The effects of disturbances

#### Ecosystem disturbance

In reaction to increasing levels of consumer mortality, the system with AF evolution behaved as the system with fixed RF. Indeed, after each increment of mortality the new biomass of consumers was similar; and the consumers disappeared for the same mortality rate (Figure 4a). Moreover, at each mortality increase, consumers in the system with AF evolution gradually reduced their foraging trait, until AF ultimately disappeared (color scale in Figure4a). Controlled monomorphic systems having low AF values better tolerated higher mortality rates (Figure 4b), which indicates that when AF was fixed it had a negative effect on the persistence of consumers facing increases in mortality.

**Figure 4:**
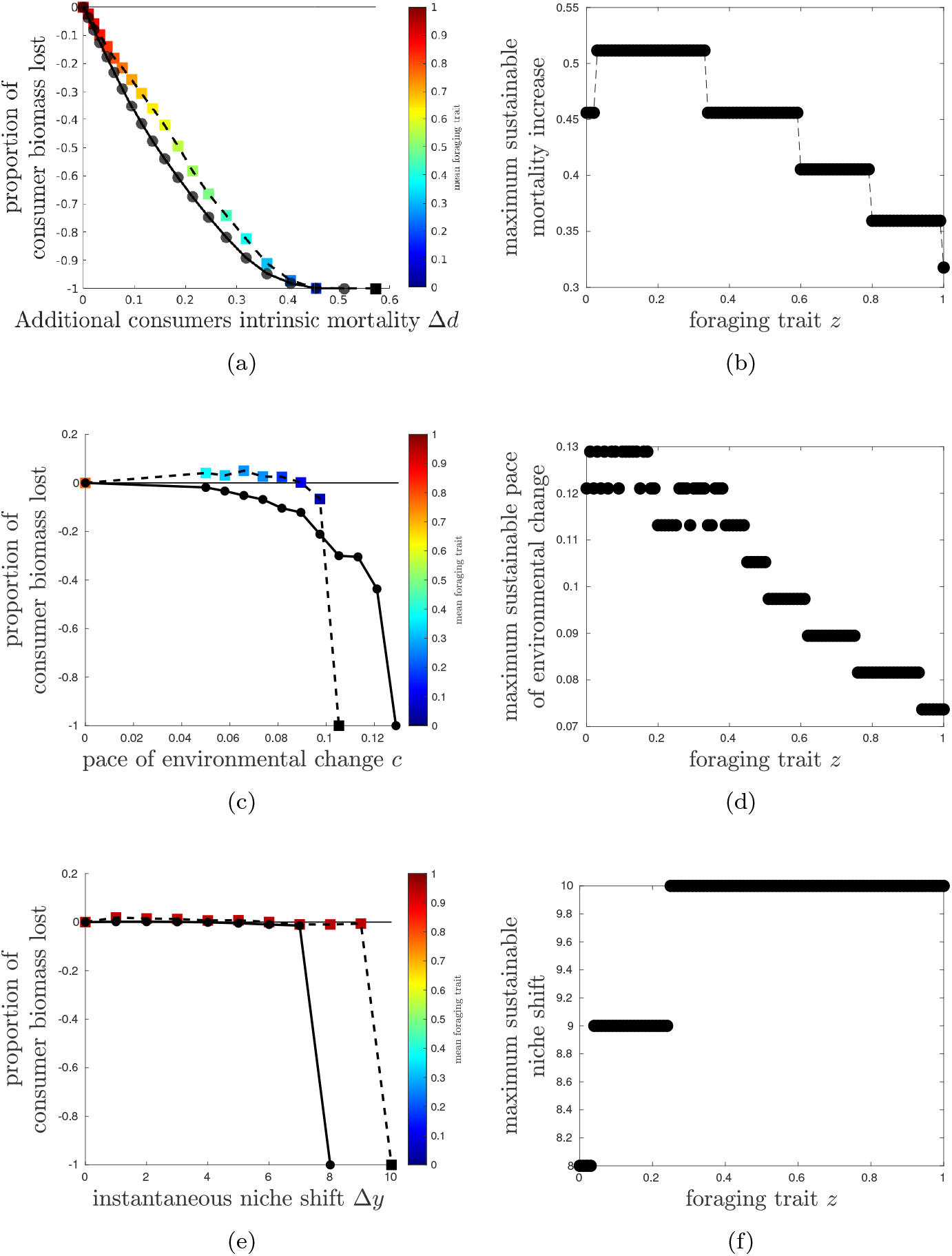
Effect of disturbances: (a, b) increased mortality ∆*d*, (c, d) constant environment chanhe *c* and (e, f) instantaneous niche shift ∆*y*. Left column (a, c, e): variations of consumer biomass of systems with and without AF, in function of the intensity of the disturbance. A negative variation indicates a decrease in biomass, for instance *−* 0.2 indicates than 20% of the biomass is lost. The value*−* 1 corresponds to the extinction of all consumers. The coloured gradient indicates the average AF trait of the consumer species. Right column (b, d f): maximal sustainable mortality for monomorphic consumers, in function of their controlled foraging trait *z*.

#### Constant environmental change

The system with AF evolution tolerated the constant environmental change better than the system with fixed RF, up to a certain point when it disappeared suddenly, earlier than its counterpart (Figure 4c). Moreover, as in the case of ecosystem disturbance, the mean AF value decreased for faster environmental changes (color scale in Figure 4c). Controlled monomorphic systems having low AF values tolerated faster environmental changes (Figure 4d), which indicates that when AF was fixed it had a negative effect on the persistence of consumers facing constant environmental change.

#### Sudden environmental change

After a sudden environmental change, either consumers disappeared or they persisted in a new state close to the original one. In that case their niche traits shifted towards the new optimum and their foraging trait remained unchanged, which is an indication of resilience. The variation of biomass before and after disturbance is therefore uninformative; instead the maximal sudden environmental change that the consumer can tolerate was used to quantify its stability (Figure 4e). The system with AF evolution resisted to a larger sudden change (*δ*_*y*_ = 10) compared with the system with fixed RF (*δ*_*y*_ = 8). In order to disentangle the direct effect of AF on stability from its indirect effect through diversity, the AF values of the consumers with AF were set to 0, while retaining the original diversity of the niche traits *x* and *y* of both guilds. The resulting hybrid system tolerated a large environmental change (*δ*_*y*_ = 10), which indicates that the positive effect of AF on the persistence of consumers was mainly due to its effects on diversity. In line with the above results, controlled monomorphic systems having high AF values tolerated larger sudden environmental changes (Figure 4d).

## 4 Discussion

### The evolutionary dynamics of adaptive foraging

Previous models exploring the effect of AF on community dynamics assumed that AF was a fixed trait of equal intensity for all consumers (Kondoh, 2003; Uchida and Drossel, 2007; Beckerman et al., 2010; Heckmann et al., 2012; Valdovinos et al., 2013). In these models, the foraging efforts of consumers changed in function of the availability and suitability of their resources, but whether foraging efforts could change or not was itself not subject to evolution. Egas et al. (2005) modelled the evolutionary dynamics of the consumers’ niche width, but not of their foraging selectivity. Therefore, the first motivation of this study was to explore under which conditions the capacity to forage adaptatively can evolve by natural selection (Question 1 in the introduction).

As expected, elevated costs of AF (*S*_*max*_, Table 1) disfavored its evolution, which is in accordance with the existence of a trade-off between AF and other life-history traits like predation (Pangle et al., 2012; Wang et al., 2013; McArthur et al., 2014; Costa et al., 2019), thermoregulation (du Plessis et al., 2012; Van de Ven et al., 2019) and time budget (Randolph and Cameron, 2001; Fortin et al., 2004). In the present model the trade-off was only incorporated into the handling time of the type II functional response, where high handling times reduced resource absorption rates. We are nevertheless confident that similar results would have been obtained if the trade-off had concerned mortality.

The evolution of AF was instead favored by the niche width of consumers (parameter *σ*). In nature, a positive correlation between total niche width and inter-individual niche variation was found for herbivores (Bison et al., 2015) and predators (Bolnick et al., 2007). Inter-individual niche variation reflects the existence of contrasting foraging strategies, which may be the result of adaptive foraging. Baboons also combine niche breadth with selectivity in resource use (Whiten et al., 1991). Since the evolution of consumer niche width may itself depend on environmental heterogeneity (Kassen, 2002) (i.e. on resource diversity in the model), the coevolution of AF, niche width and niche position is a possible avenue for future research. Niche width fostered AF because consumers depleted the whole range of resources when their niche width was large, therefore competition between consumers was more intense, which led to the evolution of AF. Empirical studies have indeed found that generalist consumers competing for resources forage adaptatively. For instance generalist bumblebee species visited the larkspur *Delphinium barbeyi* when the most abundant bumblebee species was experimentally removed, but preferred other plant species otherwise, likely to avoid competition for nectar (Brosi and Briggs, 2013). A similar behavior has been reported for syrphid flies, which preferentially foraged on open rather than tubular flowers when competing with bumblebees (Fontaine et al., 2006). In the case of predators, intraspecific competition between sticklebacks (*Gasterosteus aculeatus)* enhanced the diversity of foraging behaviors and increased the correlation between diet and morphology (Svanbäck and Bolnick, 2007), as found here (Figure A4).

The present model further predicted that AF evolution is favoured by direct competition between consumers *I* (other than for resources) as well as by increased consumer mortality *δ*. This is in line with the above results, in the sense that constrained environmental condition for consumers strengthen the need for AF. On the other hand AF becomes useful when resources are diversified enough, hence the positive effect of the resources niche width *σ*_*K*_.

The results discussed above are based on the mean foraging trait 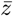 but consumers positioned at the niche edge foraged adaptatively much more often than those at the niche center (Figure 2b). Indeed, scarce resources located at the niche edge were consumed significantly by adaptive foragers only, whereas abundant resources located at the niche center could be consumed in large amounts by random foragers. This model prediction calls for empirical testing, as we are not aware of any existing work reporting this pattern.

### The effects of AF on functional diversity

Starting from a fixed pool of species or phenotypes, most previous theoretical works have shown that AF fosters food web complexity and community stability (Kondoh, 2003; Uchida and Drossel, 2007; Beckerman et al., 2010; Heckmann et al., 2012), although this depended on the way AF was incorporated to the model (Berec et al., 2010). However, had niche traits been also subject to evolution, AF might also have affected stability indirectly, through its effect on functional diversity (Figure 1). The effects of AF on diversity and other community-level properties (Question 2 in the introduction) are discussed in the present section and the effects on stability (measured by consumer persistence) in the following section (Question 3).

As expected, the evolution of AF decreased niche overlap between consumers (Figure 3d). AF also decreased niche overlap between pollinators in the model of Valdovinos et al. (2013) and in the experiments discussed above (Fontaine et al., 2006; Brosi and Briggs, 2013). At the intraspecific level, niche overlap between individuals of the same species decreases in function of their abundance (Svanbäck and Bolnick, 2007; Tur et al., 2014), suggesting the existence of a plastic behavior. Since abundance favors intraspecific competition, this is consistent with our findings that competition between consumers promotes the evolution of AF. The decrease of niche overlap between consumers corresponds to niche partitioning, which may favor their coexistence (Behmer and Joern, 2008; Turcotte and Levine, 2016). Indeed, in the model the evolution of AF enhanced the functional diversity of both consumers and resources (Figure 3b), due to an eco-evolutionary loop between resources and consumers situated at the niche edge. Following the evolution of AF some consumers foraged at the niche edge, thereby reducing the density of the corresponding resources. This decreased competition among these resources and promoted the emergence of new resource phenotypes at the niche edge. The diversification of resources triggered the apparition of consumers standing even further away from the niche centre, and so on until the resources reached the limits of the exploitable niche. This emphasizes that adaptive phenotypic plasticity like AF can subsequently fuel evolutionary change (Baldwin, 1896; Crispo, 2007; Laland et al., 2014). Instead, when no AF evolution was introduced, the few resources standing far away from the niche centre were barely used by consumers, which could not forage preferentially on them. This prevented the emergence of new resources further away from the niche centre, due to competition between resources. Since the evolution of AF occurred when the diversity of resources was initially large enough (large *σ*_*K*_), causation was reciprocal: AF both promoted and was promoted by resource diversity.

Following the evolution of AF, the functional complementarity and diversity of consumers increased their biomass at the expense of resources (Figure 3a). This fits with empirical studies showing a relationship between resource consumption and consumer diversity (Deraison et al., 2015; Lefcheck et al., 2019; Milotić et al., 2019). The effects of AF on productivity, defined as the net production of biomass by consumers following resource absorption (Table A1) were instead contrasted (Figure 3c). Moderate values of AF 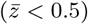increased productivity thanks to functional complementarity between consumers (Poisot et al., 2013), but higher AF values decreased productivity because consumers impacted resources too heavily.

### The effects of AF on consumer persistence

After a sudden environmental change corresponding to an instantaneous shift of the niche center, consumers with AF evolution withstood larger disturbances (Figure 4e). Previous theoretical studies have shown that AF can stabilize foodwebs by favoring more robust topologies able to buffer environmental disturbances (Kondoh, 2003; Heckmann et al., 2012). In the present model, the mechanisms responsible for this observation also rely on the dynamical nature of the interaction webs produced by AF, but not on the emergence of robust topologies. One of these mechanisms is caused by a direct effect of AF (Question 3a), and the other by an indirect effect through diversity (Question 3b), as detailed in the results. The direct effect of AF on consumer persistence relies on the mitigation of the lag load faced by consumers. Indeed, resources became adapted to the new niche center more quickly than consumers, which suffered from a trait mismatch (e.g. Post and Forchhammer, 2008; Miller-Struttmann et al., 2015; Damien and Tougeron, 2019). This indicates that phenotypic plasticity acted as a rapid response mechanism to environmental change (Fox et al., 2019), in that case. Since random foragers consumed the most abundant resources (but not the most suitable), after a sudden niche shift they fed on sub-optimal resources, which hampered their resilience to environmental change. In contrast adaptive foragers selected less abundant but more suitable resources, which favored their survival. In the meantime their traits evolved towards the new niche optimum and ultimately caught up the resources, which illustrates that adaptive plasticity can promote persistence in new environmental conditions (Ghalambor et al., 2007).

Turning to the indirect effect of AF on consumer persistence (Question 3b), when AF increased the diversity of both resources and consumers this favored the emergence of extreme phenotypes far away from the niche center. The extreme phenotypes were pre-adapted to the niche shift and therefore persisted, unlike the central species. The positive effect of biodiversity on ecosystem functioning can be caused by complementarity and selection effects (e.g. Loreau and Hector, 2001). In the present case, a few well-adapted phenotypes determined the resilience to the niche shift : this corresponds to a selection effect. Although AF also increased complementarity between species as discussed earlier, this did not created any synergy between phenotypes, at least with respect to the resilience to the niche shift.

In the cases of ecosystem disturbance and constant environmental change, AF had this time a negative effect on consumer persistence, as indicated by simulations with fixed AF values (Figures 4 b and d). For both disturbances the cost of AF became larger than the benefits, and choosy consumers went extinct earlier than random consumers. In particular, constant environmental changes weathered resource diversity to such a point that RF and AF consumers had a similar diet, which annihilated the benefits of AF. It has been stressed that phenotypic plasticity can retard adaptation to environmental change, shielding suboptimal phenotypes from natural selection (Fox et al., 2019), but in the present model phenotypic plasticity was limited to the foraging strategy of consumers. Instead, niche traits were not plastic and were therefore entirely sensitive to selection; the negative effect of AF on consumer persistence was therefore only due to its cost. In nature however, niche trait can also be plastic (e.g. Rossiter, 1987), but this was ignored by the model.

In figures 4 b and d AF was fixed but when AF could evolve, it gradually decreased in function of the intensity of the disturbances (see color scales in Figures 4 a and c). In the case of a particularly fast environmental change, consumers did not have enough time to reduce their AF searching behaviour and became extinct slightly earlier (Figure 4c). The constant environmental change created a lag load to consumers, whose niche traits ran after those of resources; in addition AF imposed a second lag load, corresponding to the time needed for the evolutionary regression of AF. In the case of ecosystem disturbance, however, since optimal foragers quickly turned into random foragers, both types of foraging strategies responded in a similar way (Figure 4a). A purely ecological model ignoring the evolutionary dynamics of AF would have missed the possibility of its evolutionary regression, and would have therefore overestimated the negative effect of AF on consumer persistence. In the simulations, the various disturbance types have been applied independently, but in nature they can be combined. In such cases, ecosystem disturbance and/or constant environmental change might first lead to the evolutionary regression of the AF behaviour, and a sudden shift might then facilitate the extinction of consumers, since they would not be protected by AF any more.

In summary, consumer persistence was fostered either by the evolution of AF in the case of a sudden environmental change or by its regression in the cases of ecosystem disturbance and constant environmental change. This corresponds to a combination of evolutionary rescue (Gonzalez et al., 2013; Kopp and Matuszewski, 2014), because AF was subject to evolution, and of plastic rescue (Kovach-Orr and Fussmann, 2013), since AF is a type of phenotypic plasticity.

### Assumption and limitations of the model

As outlined earlier, compared with other existing models exploring the influence of AF on community stability, the main novelty of the model is to study the evolution of the propensity to forage adaptatively, together with the evolution of niche traits of resources and consumers. Several other specificities also require some consideration.

First, in previous works the absence of AF corresponded to a constant interaction matrix between resources and consumers (e.g. Kondoh, 2003; Valdovinos et al., 2013). Instead, in the present model the alternative to adaptive foraging consists in random foraging, where resources are consumed according to their density. The interaction matrix is therefore highly dynamic for both foraging strategies, although for different reasons. In the case of RF the resources exploited by a given consumer change according to their abundance only, whereas in the case of AF they also change according to their traits, the consumer’s trait, and their degree of exploitation by other consumers. In previous models allowing the evolutionary diversification of niche traits, the interaction matrices were dynamic but consumers did not forage adaptatively (Loeuille and Loreau, 2005; Allhoff et al., 2015). In those cases as well as here, new phenotypes constantly appear and need to be incorporated into the food web, which is therefore inherently dynamic (Appendix A.2). In comparison to RF, a consumer having fixed interaction coefficients would ignore these new phenotypes even if its favorite resources had gone extinct, which would make little sense. Besides, AF alone can produce non-equilibrium dynamics even with a fixed community composition, by triggering consumer-resource cycles (Abrams, 1992; Abrams and Matsuda, 2004).

Second, it was assumed that consumers feeding on a single optimal resource had the highest growth rate. Although this assumption often fits with prey-predator interactions (but see Jensen et al., 2012, for a counterexample), in the case of plant-herbivore interactions consumers often benefit from resource complementarity (Abrams, 2010; Unsicker et al., 2008), primarily because of nutrient balancing and toxin dilution (Ibanez et al., 2012; Behmer and Joern, 2008; Singer et al., 2002). We predict that the inclusion of this feature in the model would have favored the evolution of AF, since RF strategists mostly consume the most abundant resources, irrespective of their complementarity.

Third, foraging costs (quantified by the searching time *s*(*z*)) were assumed independent of resource abundance, although the searching time may be larger for rare than for abundant resources. Moreover, the spatial distribution of resources were ignored, although travel time is costly (WallisDeVries, 1996; Hassell and Southwood, 1978). For instance, the random distribution of low preferred plant species can disfavor herbivore foraging selectivity (Wang et al., 2010). These two factors may hamper the evolution of AF.

## Conclusion

The present model illustrates how phenotypic plasticity can be simultaneously a result and a factor of evolution. On the one hand, adaptive foraging (AF) evolved by natural selection acting on consumers. On the other hand, it stimulated the diversification of ecological characters not only of consumers but also of resources, stressing that phenotypic plasticity can have far-reaching evolutionary consequences at the community-level (Fordyce, 2006). Moreover, functional diversity itself promoted the evolution of AF, creating an eco-evolutionary feedback loop between phenotypic plasticity, natural selection and community composition. This had intricate consequences on the response of the resource-consumer community to disturbances. In the case of sudden environmental change, the evolution of AF had a positive effect on community stability, partly via its effects on functional diversity. However for other disturbance types like constant change and increases in mortality, the AF behavior was less fit than random foraging and therefore declined. In contrast to previous studies, these results stress that the relationship between AF and community stability depends on the type of the disturbance as well as on the evolutionary dynamics of AF itself.

## Author contributions

SI, JG and LL originally formulated the project; SI, JG, EF and LL developed the model; LL and OG performed the numerical analyses; all authors participated in writing the manuscript

## Appendix

### A.1 Model details

**Figure A1:**
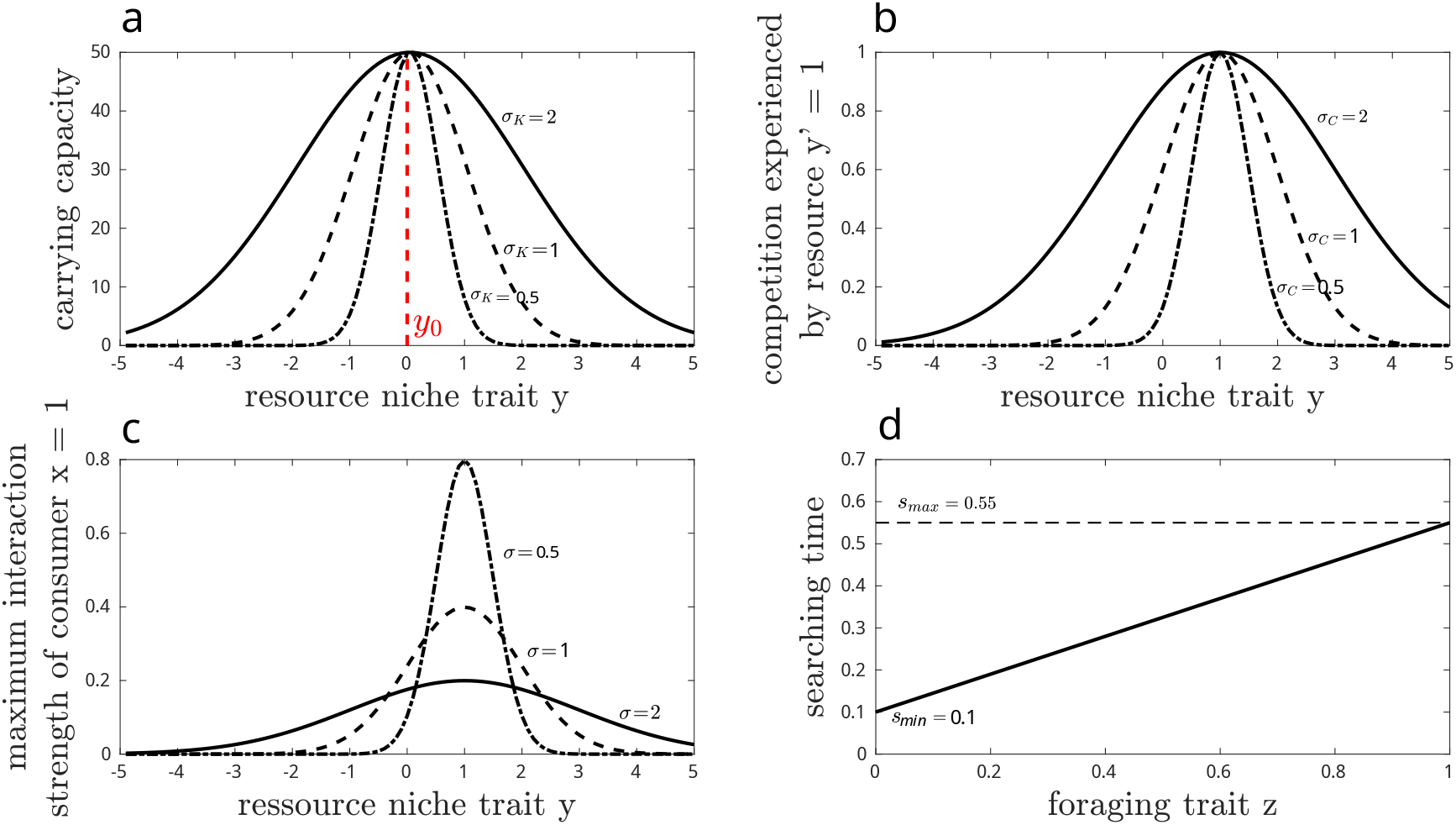
a) Carrying capacity *K*(*y*) of resources for various niche width values *σ*_*K*_ = 0.5, 1, 2. The niche centre fixed at *y*_0_ = 0 corresponds to the maximal carrying capacity. b) Competition kernel *K*_*eff*_ for various neighbourhood size *σ*_*C*_ = 0.5, 1, 2 between a focal resource *y* = 1 and all resources in function of their niche trait *y*. c) Interactions kernel ∆ for various generalization levels (*σ* = 0.5, 1, 2) between a focal consumer (*x* = 1) and all the resources in function of their niche trait *y*. d) Searching time *s* in function of the foraging trait *z*. Parameter values as in Table 1.

**Table A1:**
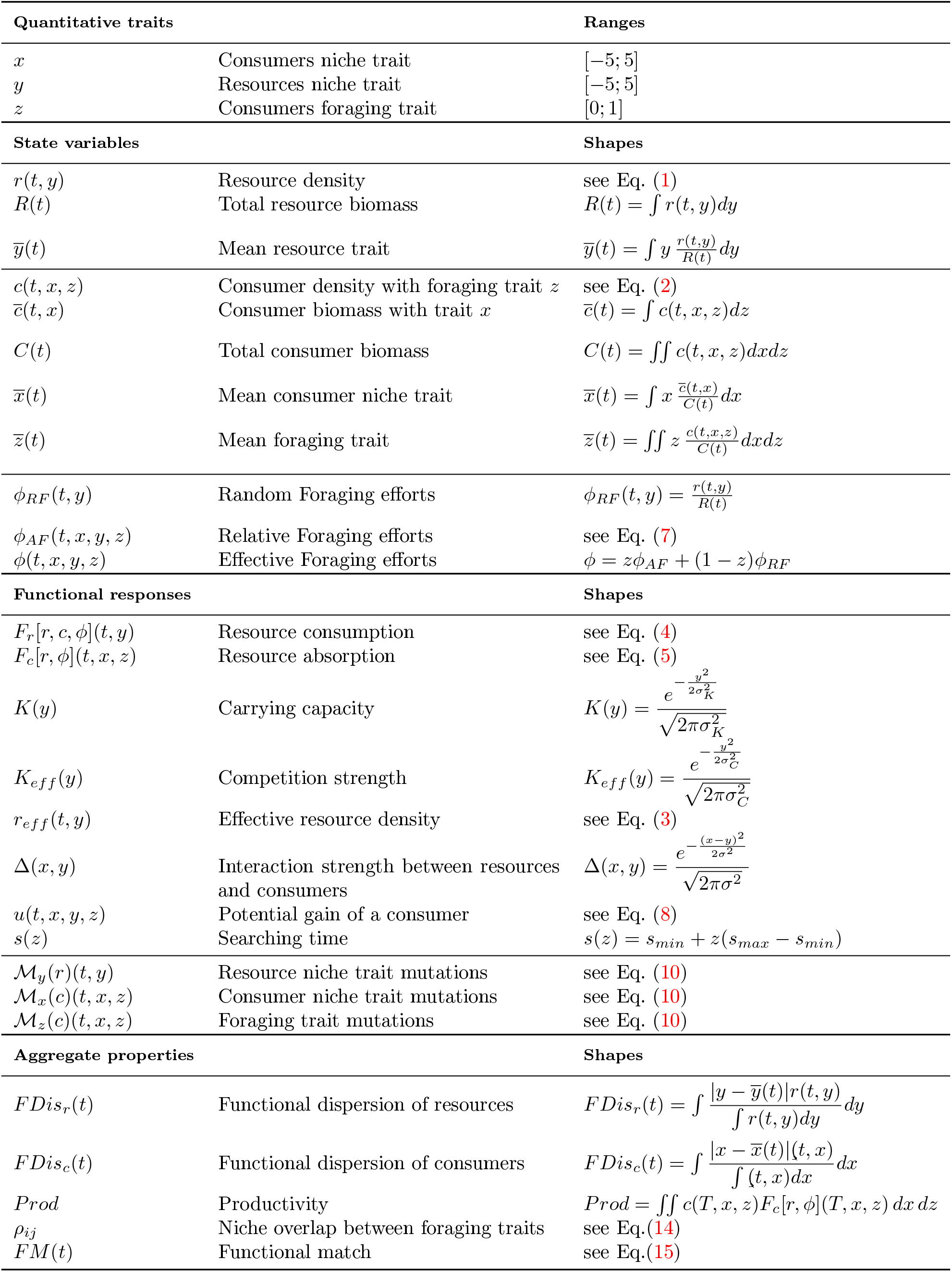
List of the quantitative traits subject to evolutionary change, the state variables, the functions and the aggregate system-level properties involved the model.

### A.2 Stationary regime

The stationary regime is visible in this simulation of the emergence of a community in which adaptive foraging evolves: https://drive.google.com/file/d/1c1nNXJl9aR76FrwFcrJppJbk-Rg7o9tn/view. The system follows a perpetual turnover of resources and consumers densities in function of their niche and foraging traits, but the macroscopic criteria of the community (exemplified here by the functional diversity *FDis*) reach a quasi equilibrium. Top panels: distribution of resources and consumers in function of their niche trait. Middle panels: distribution of consumers in function of their foraging trait (left) and community-level mean foraging trait in function of time (right). Bottom panels: functional diversity *FDis* of resources and consumers. The other community-level characteristics are also stabilized once the stationary regime is reached.

### A.3 Effect of a fixed AF trait

**Figure A3:**
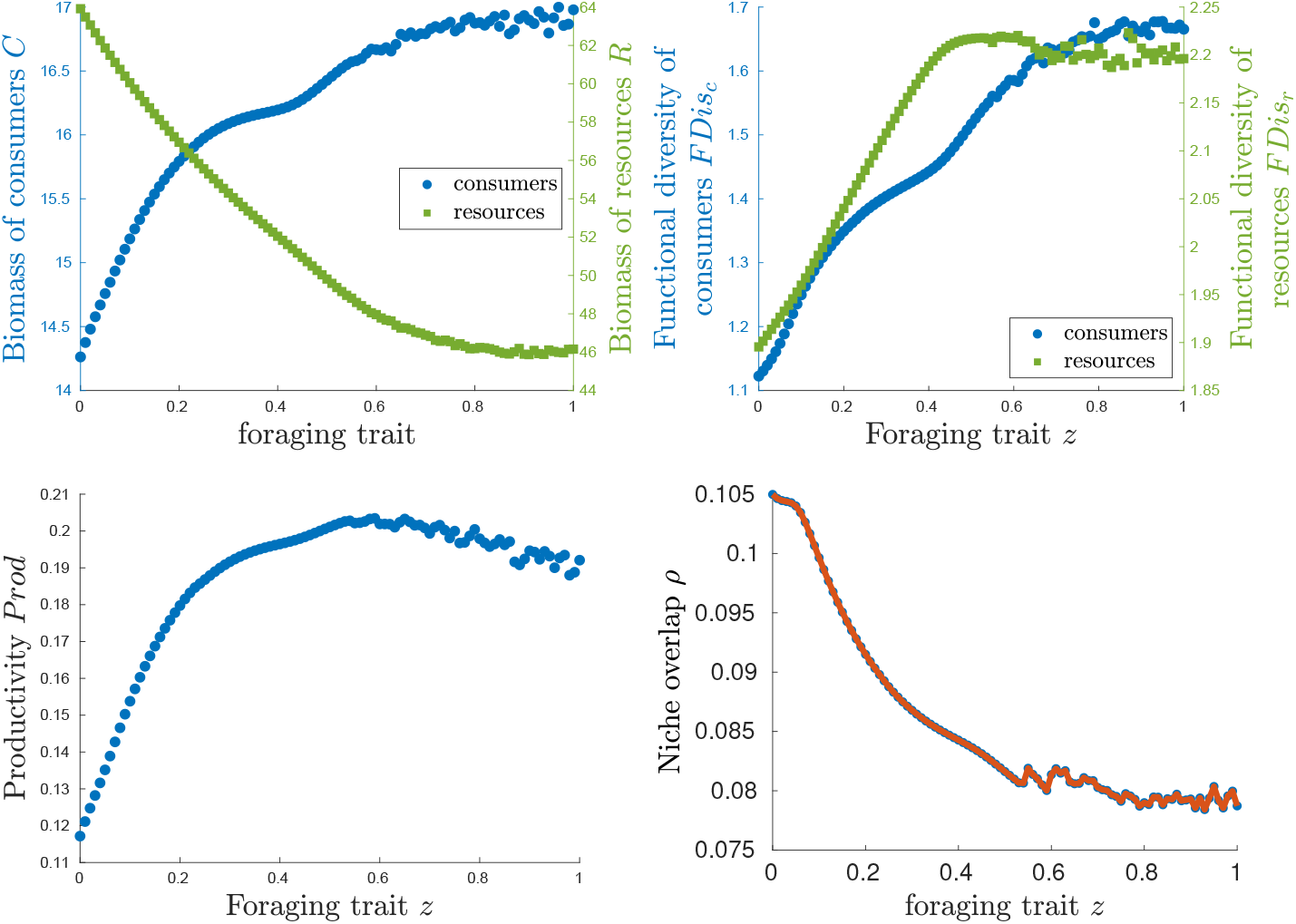
Effect of a fixed foraging trait value *z* on systems where only the niche traits *x* and *y* of resources and consumers can evolve. The measured characteristics are biomass, functional diversity, productivity, and niche overlap.

### A.4 Functional match between resources and consumers

**Figure A4:**
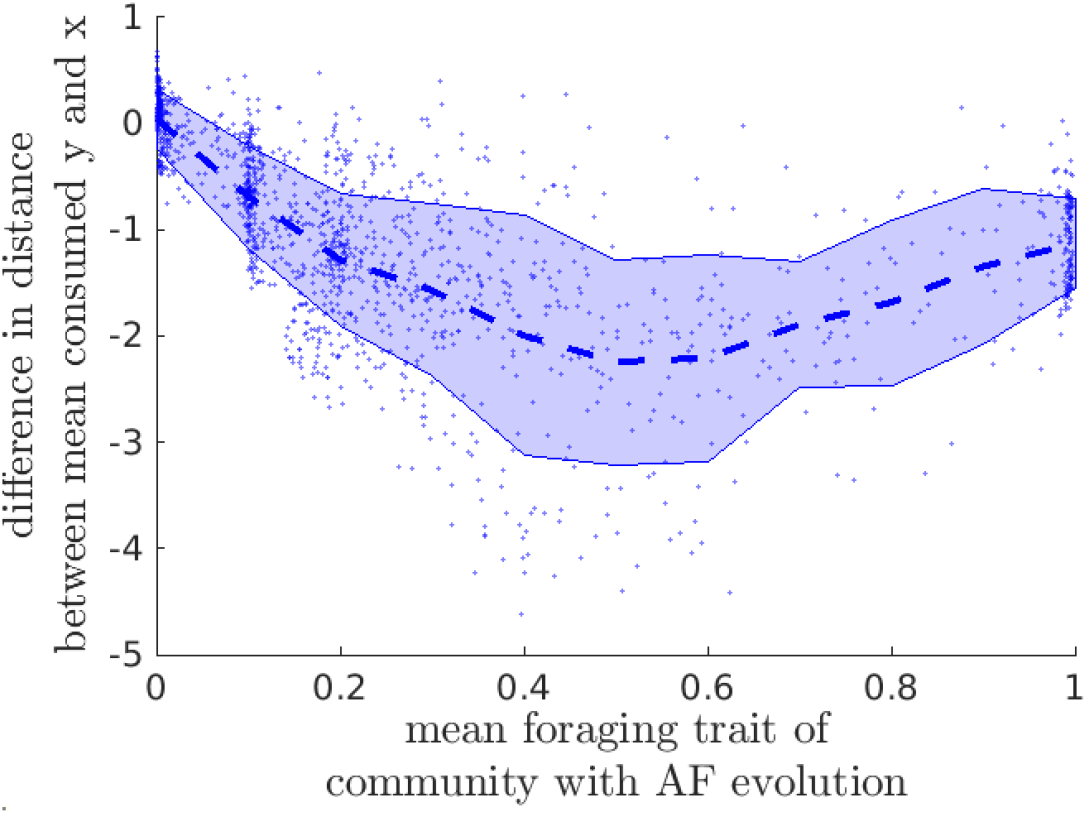
Difference in functional matching between systems with AF evolution and systems with fixed RF. 500 pairs of systems were compared, each pair having the same parameter set randomly sampled in the ranges specified in Table 1. Dashed lines: median; areas: 75% confidence intervals.

